# Massively Concurrent Sub-Cellular Traction Force Videography enabled by Single-Pixel Optical Tracers (SPOTs)

**DOI:** 10.1101/2023.07.25.550454

**Authors:** Xing Haw Marvin Tan, Yijie Wang, Xiongfeng Zhu, Felipe Nanni Mendes, Pei-Shan Chung, Yu Ting Chow, Tianxing Man, Hsin Lan, Yen-Ju Lin, Xiang Zhang, Xiaohe Zhang, Thang Nguyen, Reza Ardehali, Michael A. Teitell, Arjun Deb, Pei-Yu Chiou

## Abstract

We report a large field-of-view and high-speed videography platform for measuring the sub-cellular traction forces of more than 10,000 biological cells over 13mm^2^ at 83 frames per second. Our Single-Pixel Optical Tracers (SPOT) tool uses 2-dimensional diffraction gratings embedded into a soft substrate to convert cells’ mechanical traction stress into optical colors detectable by a video camera. The platform measures the sub-cellular traction forces of diverse cell types, including tightly connected tissue sheets and near isolated cells. We used this platform to explore the mechanical wave propagation in a tightly connected sheet of Neonatal Rat Ventricular Myocytes (NRVMs) and discovered that the activation time of some tissue regions are heterogeneous from the overall spiral wave behavior of the cardiac wave.

**One-Sentence Summary:** An optical platform for fast, concurrent measurements of cell mechanics at 83 frames per second, over a large area of 13mm^2^.

## Main Text

Mechanical force regulates biological processes at scales spanning molecules (*1*), cells (*2*), tissues (*3,4*), organs (*5*), and up to enabling the survival and growth of a multicellular organism by facilitating, for example, heart beating (*6*), blood vessel contraction (*7*), intestinal food digestion (*8*), and body movements (*9*). These multicellular activities require the coordination of complex combinations of electrical, chemical, and mechanical processes (*10*). Tools and techniques for analyzing multicellular systems are readily available in the electrical (*11*), and chemical domains (*12*), but approaches for assessing multiscale mechanical processes are currently lacking and inferred. This technology gap is especially evident for rapid and dynamically coordinated events that can involve the collective interactions of many cells over large distances (*13*).

Current approaches used to quantify mechanical properties of cells and tissues include atomic force microscopy (AFM) (*14,15*), traction force microscopy (TFM) (*16*), elastic pillars (EPs) (*17,18*), muscular thin film (MTF) (*19*), and fluorescently labelled elastomeric contractible surfaces (FLECS) (*20*). AFM provides high spatial resolution and high sensitivity measurements but depends on probe tip placement and is low throughput (*21*).

TFM quantifies traction forces by tracking the movement of nanoparticles embedded in an elastic substrate (*16,22,23*). EPs extract cellular traction forces by measuring the bending of elastic micropillars (*24*). Both TFM and EP methods require high numerical aperture (N.A.) optics to determine nanoparticle or micropillar movements for accurate measurements. The small field-of-view (FOV) of high N.A. optics, however, limits the number of cells for concurrent assessments, which is the major hurdle for studying interconnected multicellular systems that change dynamically during short time periods. Measurement accuracy and reproducibility may be adversely affected when data from numerous cells in a large area is generated by assembling sequential readings taken from smaller regions within a more extensive process, particularly when the conditions of the cells change during the measurement period. Elastomer post and MTF approaches measure the average force output of a group of cells based on the deformation of a pillar or a film. These force averaging methods lack information on individual cells, subgroups of cells, and their synergistic or antagonistic effects. FLECS can provide high throughput single cell measurements, although this approach requires seeding isolated cells on tracer structures, thereby preventing FLECS from quantifying interconnected multicellular systems.

Current measurements of mechanical wave propagation in tissues use optical mapping (OM) methods (*25-28*). However, OM has several fundamental limitations. First, OM requires cell labeling with fluorescing dyes, such as Di-8-ANEPPS or Fluo-4 AM. Compromised cell viability can occur with OM because of photochemical toxicity related to high concentration dye and intense illumination for high-speed fluorescence imaging over extended imaging times, and probe quenching can limit periods of data collection. Second, OM is an indirect force measurement method. It relies upon cells’ electrical or chemical signals to indicate cells’ mechanical responses. The relationship between these indirect indicators and mechanical activity and strength are inconsistent and vary amongst cells (*25-28*). Furthermore, fluorescent detection does not provide information on force direction, which is a crucial parameter for constructing traction force distribution. As a result, it is not possible to extract traction force data using traditional OM methods.

Here, we introduce an innovative concept and approach called Single-Pixel Optical Tracers (SPOTs) microscopy. SPOT microscopy directly measures rapid coordinated or asynchronous changes in traction forces of cells within a large interconnected multicellular sheet with sub-cellular resolution. This approach overcomes major limitations of conventional OM because there are no fluorescing labels, eliminating photochemical toxicity and probe quenching over extended imaging times. SPOT microscopy is also a direct force measurement method. Color changes recorded from SPOT images emerge from grated micromirror tilting that is proportional to applied traction forces coming from overlying cells. This feature removes the uncertainty of indirect mechanical response measurements inferred from electrical or chemical signaling methods.

## Results and Discussion

SPOT microscopy requires a platform of periodic, arrayed micromirrors that harbor micro-fabricated reflective optical gratings (Fig. 1a, fig. S1). Under broadband light illumination, each micromirror grating separates the light into its constituent frequencies, reflecting a rainbow of colored light rays. When overlying cells apply directional force on the micromirror-containing substrate, a micromirror tilts, which tilts the reflected color rainbow. A low N.A. Canon MP-E 65mm f/2.8 optical lens collects the reflected light rays from the micromirror array onto an image sensor (see Materials and Methods). The low N.A. lens functions as a bandpass color filter to permit the collection of a narrow, reflected color band (Δθ) from each micromirror in a large imaging field. When a specific micromirror in the imaging field tilts, its reflected band shifts, causing a color change on the image sensor (Fig. 1b).

**Fig. 1.**
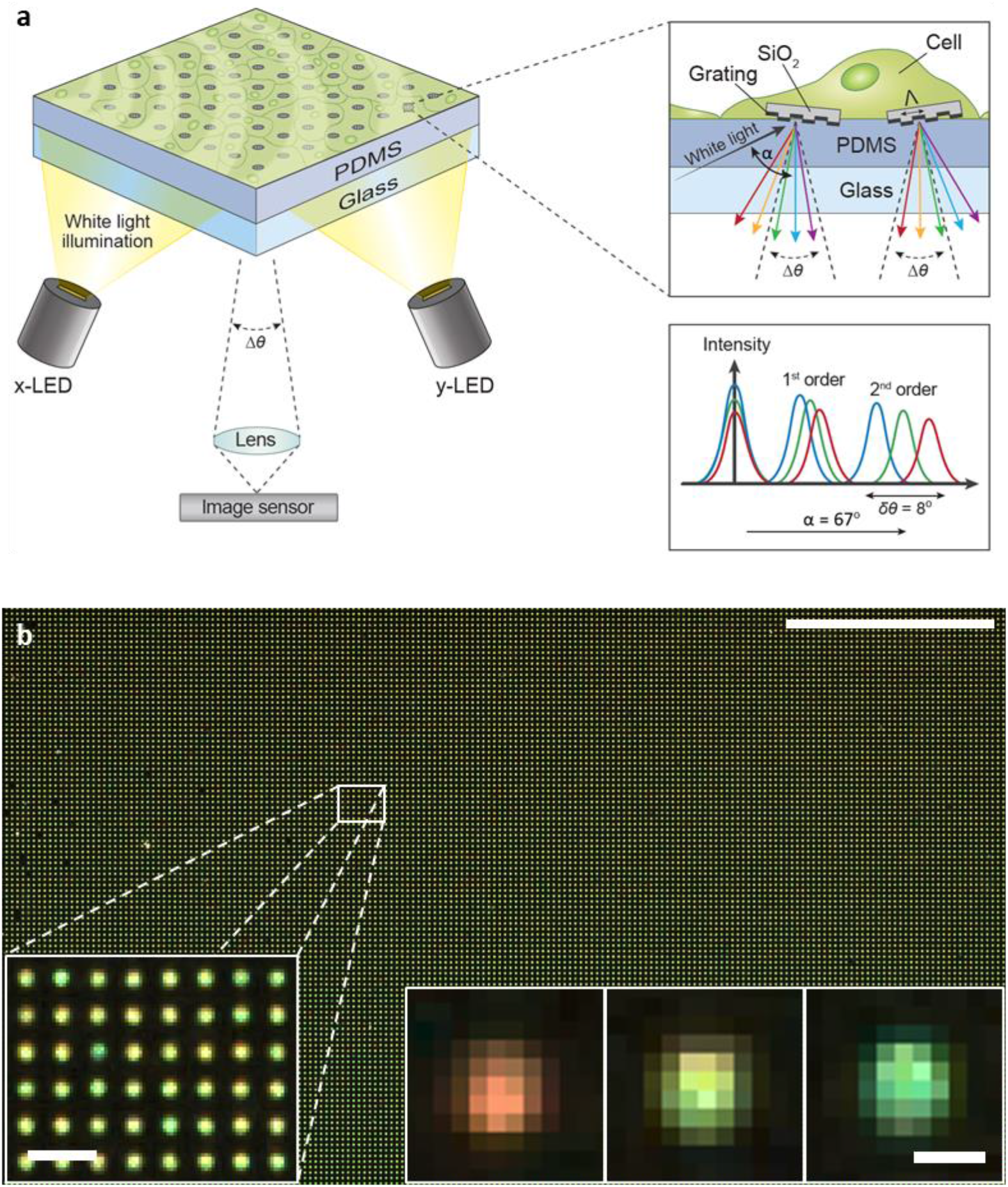
SPOT microscopy for quantitative measurements of mechanical wave propagation in two-dimensional interconnected tissues. **a**, SPOT microscopy consists of an array of diffraction grating micromirrors bonded on top of an elastic PDMS sheet. Cells exert traction force on the substrate and cause the micromirrors to tilt. Each grating micromirror reflects and disperses incident white light into microscale rainbow light rays. Second-order diffraction maxima are chosen for their optimal dynamic range and sensitivity in measurements. A low N.A. lens is used to collect a narrow color band from the rainbow light rays of each micromirror for imaging. When a micromirror is tilted, the collected color band changes according to the extent of tilting. **b**, A representative image captured by SPOT microscopy. The bottom right inset shows the micromirror images that represent tilting angles of -4, 0, and +4 degrees, respectively. (Scale bars: top right, 1 mm; bottom left, 50 µm; bottom right, 10 µm)

Quantitative measurement of traction force requires data that includes the force direction, which demands a measurement of micromirror tilting angles in both the x and y directions.

To obtain micromirror tilting angles, we performed Gaussian intensity fitting on imaging pixels for each micromirror in each of the three color channels (R, G, B). Since there are two directions, x and y, the tilting angle status of each micromirror is encoded within six color values, three for the x-direction and three for the y-direction. After obtaining the RGB values for each micromirror, we further convert the RGB color scheme into the hue-saturation-value (HSV) color scheme because the Hue value is an ideal parameter for quantifying the color band reflected by a micromirror as it represents the dominant wavelength of a color. Our current iteration of the SPOT microscopy micromirrors platform can measure tilting angle changes between -4 to +4 degrees in both x and y directions (fig. S2).

To extract the traction force distribution of Neonatal Rat Ventricular Myocytes (NRVMs) from measured micromirror tilting angles using SPOT microscopy, we applied a machine learning method to train on 2,000 simulated examples of tilting angles generated by the finite element method (FEM) using COMSOL Multiphysics. Each simulated example maps a 2D random traction force distribution to a tilting angle distribution of a 5×5 micromirror array (see Materials and Methods). The trained model calculates the traction force distribution for a small 5×5 micromirror array, and therefore we fed SPOT microscopy measured tilting angle data into this model sequentially in batches (Fig. 2a). For each calculation, we select only the traction force distribution calculated near the center micromirror for subsequent processing to avoid boundary condition effects in the simulation (Fig. 2b). We filled junction areas between neighboring micromirrors using interpolation (Fig. 2c,d). We repeated this process to extract the traction force distribution on each micromirror across an entire 4.8 mm × 2.7 mm FOV (Fig. 2e).

**Fig. 2.**
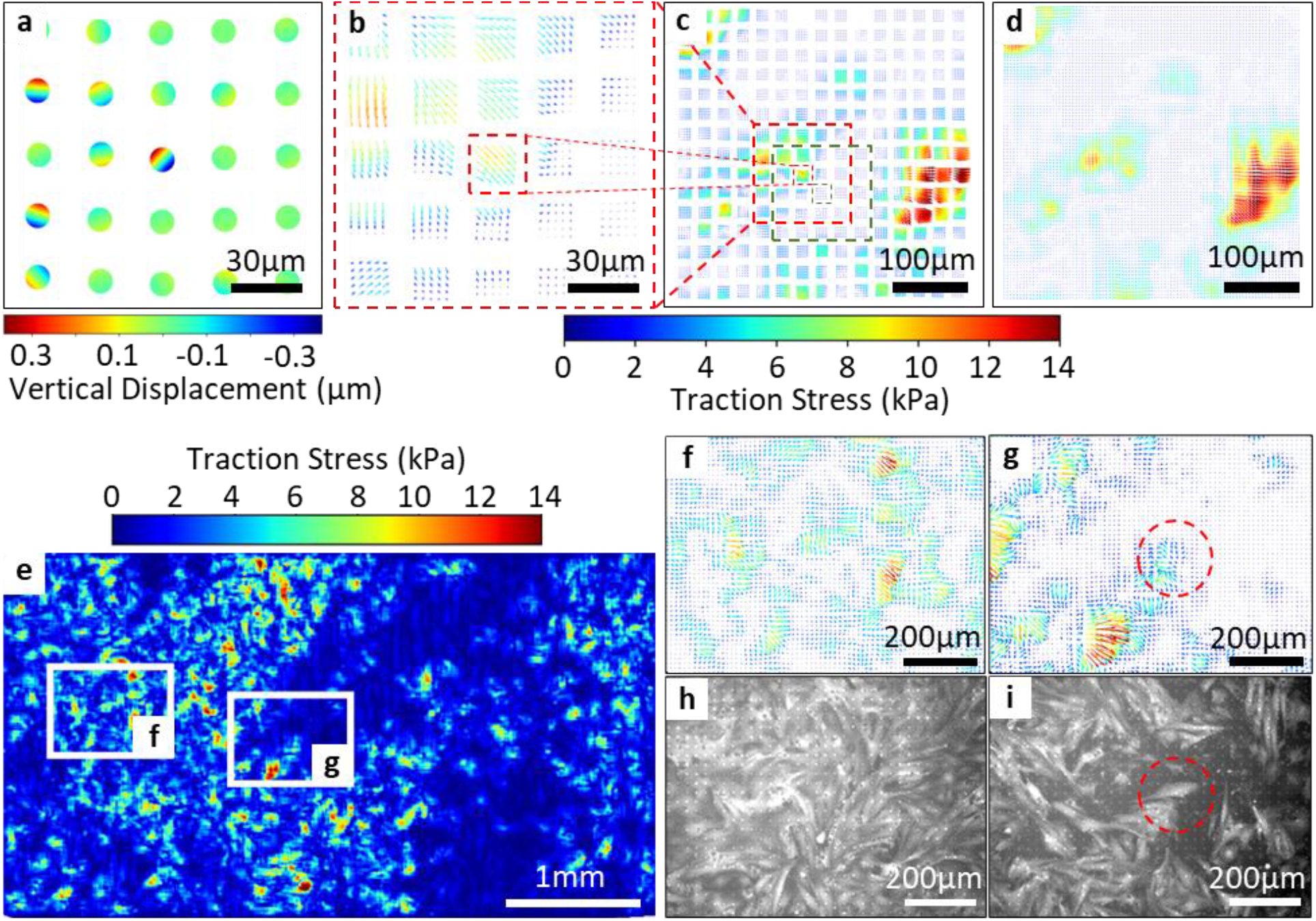
Traction force distribution measurements using SPOT microscopy. **a**, Experimentally measured tilting angles of a 5**×**5 micromirror array by SPOT microscopy. **b**, A trained machine learning model computes the traction force distribution across the 5×5 micromirror array example in (**a**). Only the traction force distribution in the vicinity of the central micromirror is utilized for subsequent construction of larger field traction force distributions to avoid edge micromirrors that show errors due to boundary condition effects. **c**,**d**, Construction of the traction force distribution for the entire FOV achieved by repeating step (**b**) and stitching through interpolation. **e**, Peak traction force distribution map of a NRVM sheet over the entire FOV. **f**,**g**, Peak traction force areas from selected regions in (**e**), with (**f**) showing a monolayer of NRVMs at 100% confluence, and (**g**) showing an area with a low density of NRVMs. **h, i**, Calcein AM fluorescent images of cells in regions corresponding to (**f**) and (**g**). The overlap between (**g**) and (**i**) at a region of nearly isolated CMs indicates that SPOT microscopy can provide sub-cellular spatial resolution of traction force distribution (see fig. S3).

SPOT microscopy records data from the entire FOV in each frame so post-data collection processing time and data stitching steps do not affect measurement accuracy. This key feature is unique from conventional small FOV methods that record data region by region at different times, which can introduce lag periods during data collection. For rapid, dynamic measurements, this is sub-optimal since cells may change significantly during sequential measurements. By concurrently measuring the dynamic tilting angles of each micromirror, we calculate the dynamic traction force for every region within the FOV. This capability enables comparisons between traction forces in different cell sheet regions occurring at different times. Consequently, we can construct a map of the distribution of maximum traction forces to quantify the beating strength of cardiomyocytes (CMs) in different NRVM regions (Fig. 2e). Magnified images of maximum traction force distributions in selected regions are shown (Fig. 2f,g), alongside corresponding fluorescence images (Fig. 2h,i) that reveal the CM distribution. Comparing these images yields an overlap between traction force distributions and cell-occupied regions. The circled areas (Fig. 2g,i) suggest that sub-cellular spatial resolution for traction force measurements is achievable using SPOT microscopy (fig. S3).

To demonstrate the unique capabilities of SPOT microscopy for quantifying mechanical wave propagation, we provide a sheet of beating, interconnected CMs as an experimental test bed. Efficient blood pumping depends on rhythmic, coordinated mechanical contraction and relaxation cycles to generate repeated, directional waves in a large sheet of physically interconnected CMs. In this context, we utilize SPOT microscopy to study two types of coordinated CM beating, the propagation of a typical linear plane wave and propagation of an atypical spiral wave, which may resemble an arrhythmia clinically. We show snapshots from a video (Fig. 3a, movie S1) of plane wave propagation using primary neonatal rat ventricular myocytes (NRVMs) imaged on the SPOT microscopy platform over a FOV of 9 mm × 9mm. The plane wave enters the imaging FOV from the right and gradually spreads over the entire CM sheet within 250ms. By 650ms, most of the NRVMs have completed at least one contraction and relaxation cycle. We also show snapshots from a video (Fig. 3b, movie S2) of a spiral wave propagation initiated at the center of the FOV that progresses in a clockwise direction. The spatiotemporal resolution of SPOT microscopy extracts dynamic intensity profiles for every pixel in the large imaging field over time. We show the time-dependent intensity profiles of four imaged pixels along the trajectory of mechanical spiral wave propagation (Fig. 3c). These profiles are direct measurements of force during contraction and relaxation cycles for individual NRVMs. SPOT microscopy can quantify multi-parametric data including activation time, contraction strength, action duration, relaxation time, beat frequency and beat pattern for each NRVM within the interconnected CM sheet over time. Heterogeneities in force generation or relaxation amongst imaged CMs may uncover patterns, trends, or relationships between individual cells, subgroups of cells, and their synergistic or antagonistic activities over large areas of measurement.

**Fig. 3.**
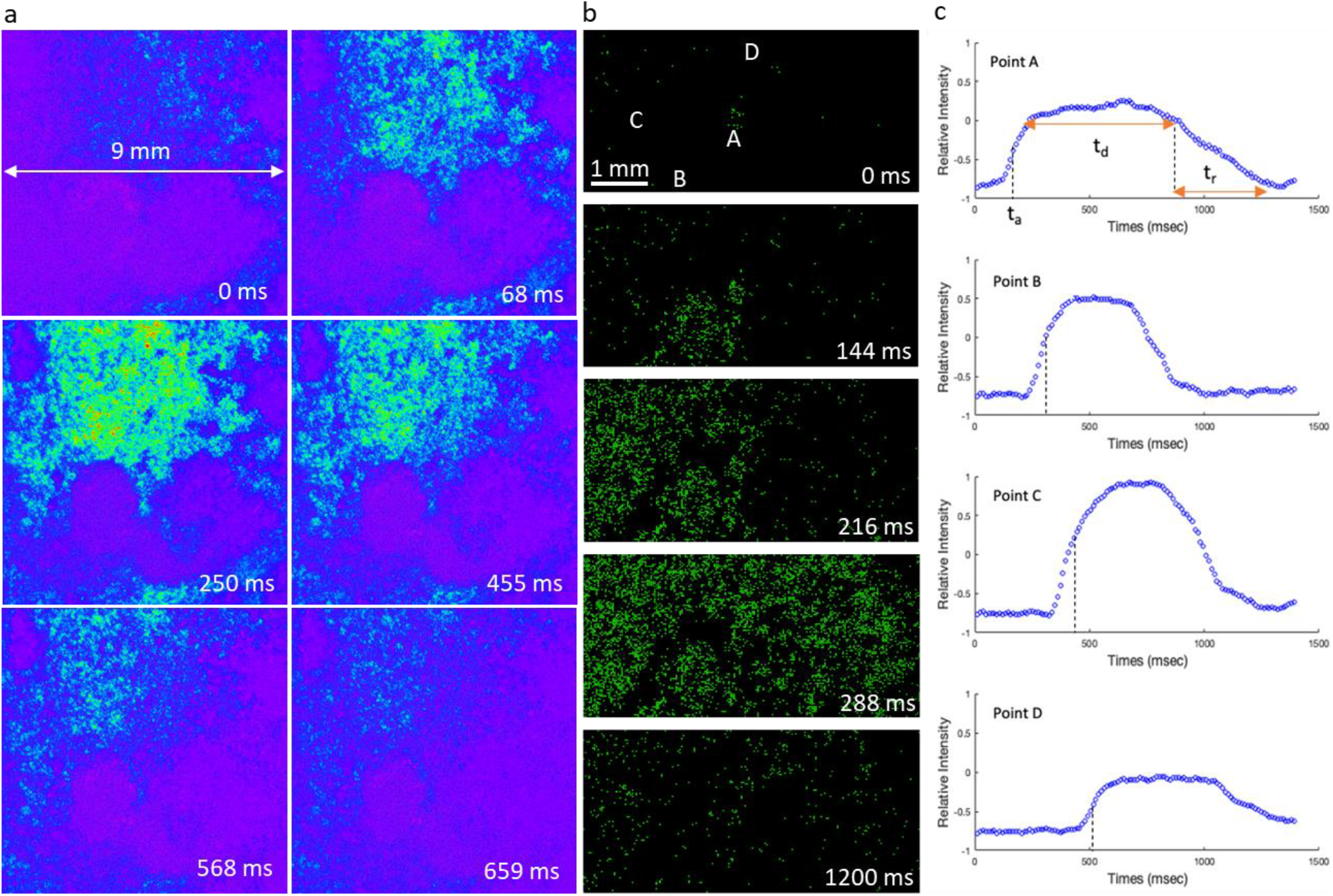
SPOT microscopy captures dynamic wave propagation of beating NRVMs. **a**, Snapshots illustrate a plane wave propagating across a 9 mm **×** 9 mm field FOV (see movie S1). **b**, Snapshots illustrate propagation of a spiral wave initiated at the center of the FOV (see movie S2). **c**, Temporal intensity profiles from selected pixels along the trajectory of a propagating spiral wave. These profiles enable the extraction of key parameters, such as activation time, contraction strength, action duration, and relaxation time, for a multi-parametric analysis of the coordinated beating behaviors of cells separated by a distance.

Application of SPOT microscopy enables the construction of an activation time map that traditionally requires OM methods to identify cells with abnormal beating rhythms. OM methods utilize fluorescent signals emitted by voltage sensitive dyes to create an activation time map. The map yields a 2D visual representation of the electrical activation sequence in cardiac tissue that uncovers the initiation, propagation and coordination of electrical signals governing the contraction and relaxation of CMs. By displaying the temporal and spatial distribution of these electrical activities, an activation time map reveals information about CM function and potential abnormalities, although different degrees of electromechanical uncoupling may yield an inaccurate representation of force generation.

In SPOT microscopy, where fluorescent signals and their experimental limitations are absent, the activation time is defined as the moment when the rate of change in the micromirror-reflected color intensity reaches its maximum. By plotting dynamic pixel intensity variations over time, we extract the activation time of each pixel from its temporal intensity profile (fig. S4). We provide the activation time map (Fig. 4a) corresponding to the plane wave propagation (Fig. 3a). This map identified heterogeneous cell groups with abnormal beating patterns and rhythms. We provide a map showing the relative contraction strength of NRVMs in different geographic regions of the CM sheet (Fig. 4b). By comparing maps of activation time and contraction strength, subgroups of cells showing distinctive contraction patterns and distributions can be identified using SPOT microscopy.

**Fig. 4.**
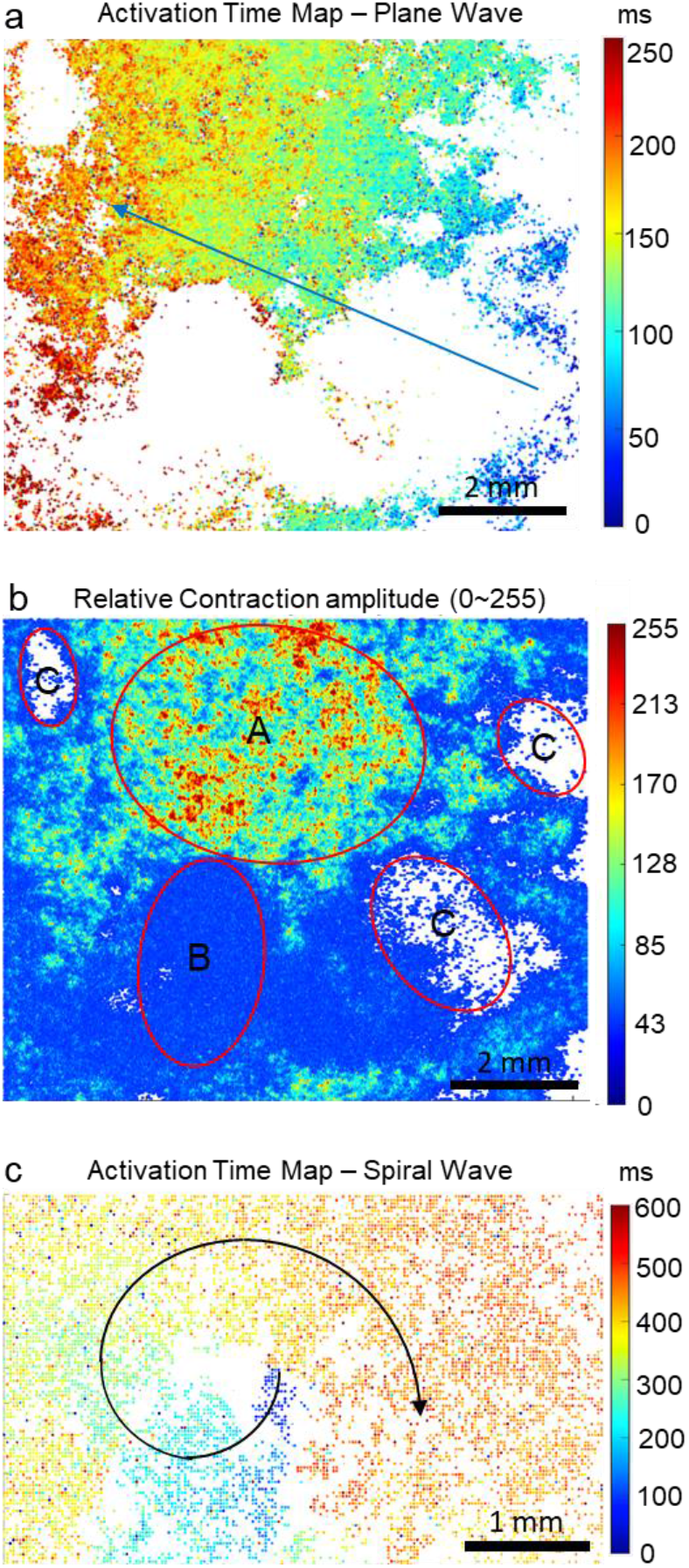
SPOT microscopy captures the heterogeneity of beating cardiomyocytes (CMs). **a**, An activation time map corresponding to the propagating plane wave in Fig. 3a. White regions represent mostly CMs with contraction amplitude below threshold and CMs whose time-domain beating data do not fit well with the logistic fitting function used for extracting the activation time (see fig. S4). **b**, A map of contraction amplitude indicates regions of distinct contraction strengths. CMs in region A show maximum contraction strength and their temporal beating patterns fit well with the logistic function used for constructing the activation time map shown in (**a**). CMs in regions B and C are groups that have contraction amplitudes lower than the threshold and were not presented in the activation time map. Yet, on the contraction amplitude plot, these two CM subgroups are clearly identified. From the bright field microscopy observation, cells in group B form a continuous sheet but have much weaker contraction strength compared to cells in group A. In contrast, C labeled regions are areas with no CMs. **c**, An activation time map corresponding to the propagating spiral wave shown in Fig. 3b. The spatiotemporal origin of the abnormal spiral mechanical wave can be clearly identified in this map as the deep blue cells at the base of the spiral arrow.

For an atypical spiral propagation wave, SPOT microscopy can identify the spatiotemporal origin of abnormal beating from a small cluster of CMs within the NRVM sheet (Fig. 4c), with potential implications for clinical translation. Cardiac arrhythmias can result from abnormal electrical spiral waves (*29*) and SPOT microscopy reveals the mechanical manifestation of this abnormal electrical activity, which connects anomalous mechanical wave propagation to irregular CM electrical excitation.

In summary, we developed a new label-free optical microscopy system to quantify rapid and dynamic mechanical wave propagation across nearly a centimeter-sized sheet of interconnected cells. Our system offers significant advantages over other indirect small FOV or fluorescence labeled approaches that suffer from potential effects on cell viability or function during measurements and limited durations of cell monitoring. In contrast, SPOT microscopy directly measures the mechanical output of cells, eliminating indirect measurements that may be variably uncoupled, and avoiding imaging-related toxicity to allow for continuous monitoring over extended periods. SPOT microscopy simultaneously records the direction and magnitude of traction forces for more than 10,000 cells, providing data with high spatiotemporal resolution. These unique multi-cellular capabilities of SPOT microscopy pave the way for a new approach and powerful platform for studies of electromechanical coupling or decoupling in CMs, arrhythmias, and drug screening, with potential for fundamental insights and ex-vivo translational applications.

## Acknowledgments

N.A.

## Author contributions

Xing Haw Marvin Tan designed and fabricated the devices, assembled, and calibrated the optical system, performed the experiments, performed the finite element simulations, trained the machine learning model, and wrote the manuscript.

Yijie Wang cultured and seeded the cells onto the devices.

Xiongfeng Zhu designed the overall fabrication process.

Pei-Shan Chung designed the fabrication process control monitoring (PCM) testing structures.

Felipe Nanni Mendes designed optical instruments using SolidWorks.

Yu Ting Chow and Tianxing Man provided technical support in maintaining the cells.

Hsin Lan, Yen-Ju Lin, and Xiang Zhang provided technical support in the experiments.

Xiaohe Zhang developed part of the machine learning code utilizing polynomial regression and B-spline regression for optical calibration.

Thang Nguyen gave technical support in the optical instrumentation.

Reza Ardehali provided neonatal rat ventricular myocytes.

Michael A. Teitell supervised the biology study and revised the manuscript.

Arjun Deb supervised the cardiomyocyte study and provided neonatal rat ventricular myocytes.

Pei-Yu Chiou proposed the concept, designed the optics, analyzed the data, and wrote the main paper.

## Data and materials availability

All data and code are available in repository Dryad which is linked to this manuscript.

## Supplementary Materials

Materials and Methods

Figs. S1 to S4

References (*01-31*)

Movies S1 to S2

