## Supplementary Materials & Methods for "Massively Concurrent Sub-Cellular Traction Force Videography enabled by Single-Pixel Optical Tracers (SPOTs)"

**The PDF file includes:**

Materials and Methods
Figs. S1 to S4

**Other Supplementary Materials for this manuscript include the following:**

Movies S1 to S2

### Materials and Methods

Optical systems

The optical setup for SPOT microscopy utilizes two high power light emitting diodes (LEDs) (Citizen CLU550-3626C1) for orthogonal illumination in x- and y-directions. The LEDs were mounted on two heat sinks to prevent overheating. Line-shaped light sources were realized by covering the LEDs with rectangular-shaped slits of 25mm×5mm dimensions. For videography at 1× magnification, a Canon EF 28mm f/1.8 Prime Lens was used with a FLIR Grasshopper3 (GS3-U3-41C6C-C) camera. For videography at 4× magnification, a Canon MP-E 65mm f/2.8 Macro lens was used with a Chronos 2.1 HD (Krontech) high speed camera. For videography at 4× magnification, the x-direction LED was switched on for 5 ms, with a 1 ms delay before the y-direction LED was switched on. The camera exposure time was 4 ms for each snapshot, yielding a frame rate of 83 fps, with each frame consisting of a pair of x- and y-direction snapshots. An Arduino circuit was used to synchronize the LEDs with the cameras.

Process flow for fabricating micromirror arrays

The fabrication process starts from a silicon wafer with a 1.56μm thermal oxide layer. The oxide layer was then patterned and etched 120nm deep to create a 2D diffraction grating pattern with linewidth and spacing of 600nm. The oxide layer was then photolithography patterned and etched to form the 10μm-diameter micromirror array. An isotropic silicon etching step was then performed to undercut the silicon under these micromirrors to leave a weak mechanical necking structure. A Ti/Al/Ti/$SiO_2$ layer was thermally evaporated onto these micromirror structure to increase the reflectivity. A separate glass wafer was then prepared and spin-coated with a 50μm PDMS layer. Oxygen plasma treatment was performed on both the PDMS and the micromirror surfaces for bonding. The micromirror array was then transfer-printed onto the PDMS surface. An ultrasound treatment was then applied to break the silicon necking structure and release the micromirror array onto the PDMS surface to finish the fabrication process. SEM imaging was used to confirm the structure of the micromirror arrays.

Two PDMS films with Young's modulus of 21 kPa (± 0.9 kPa) and 197 kPa (± 22 kPa) were fabricated for SPOT microscopy measurements. The Young's modulus of PDMS films were characterized using an Instron 5564.

Cell seeding for imaging

Dopamine (0.01%, w/v in 1M Tris-HCl buffer) was coated onto the surface of the SPOT microscopy platform at 4°C for 1h according to Chuah *et al* (*30*). Then, 35 min of ultraviolet sterilization was followed by coating with Matrigel (83 µg/mL) on the device surface for 1h. We seeded NRVMs on the SPOT microscopy platform at 50% ~ 100% cell confluency with overnight incubation at $37^oC$, 5% $CO_2$ before imaging measurements using SPOT microscopy.

Machine learning model for traction force distribution measurements

A linear regression machine learning model (scikit-learn) was trained and used to predict the traction force distribution from the micromirror tilting angle data collected using SPOT microscopy. This machine learning model used training data that includes random stress distributions as inputs and tilting angles of micromirrors calculated by the finite element method (FEM) model in COMSOL as outputs.

Random stress distributions were generated from a 2-dimensional Fourier series, using random numbers for the amplitude and phase terms (Eqn. 1):

$$Stress(x, y) = \sum_{-n}^{n} \sum_{-m}^{m} A_{m,n} \cos[w(mx + ny) + \phi] \qquad (Eqn.\ 1)$$

$$where\ w = \frac{2\pi}{\lambda} = 2\pi f$$

$$\lambda = 1.8mm$$

$$A_{m,n}\ and\ \phi\ are\ random\ numbers.$$

A total of 2,000 different random stress distributions were imported into COMSOL to calculate the corresponding outputs of tilting angles of a 5 × 5 micromirror array. These 2000 training datasets were used to train a linear regression machine learning model.

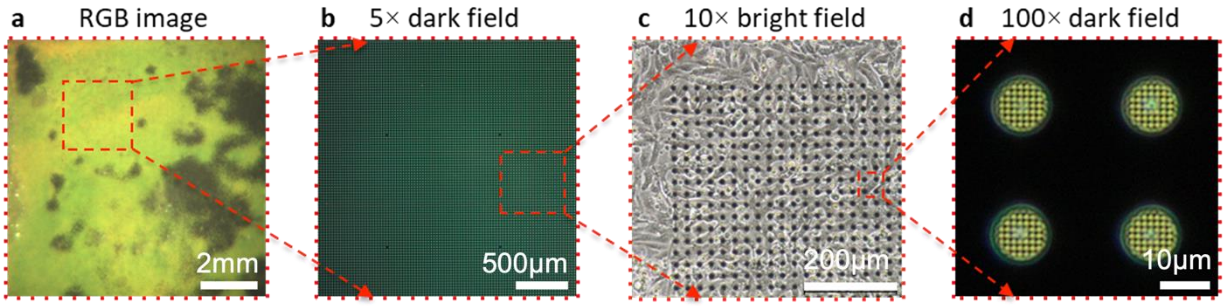


**Fig. S1.**

**Comparison of SPOT microscopy images with microscopic images captured under different magnifications.**

**a**, A whole 9mm × 9mm FOV image captured by SPOT microscopy.

**b**, A 5× dark field image showing an array of micromirrors on the SPOT microscopy platform.

**c**, A 10× bright field image showing the relative size of micromirrors and a sheet of NRVMs cultured on a SPOT microscopy platform.

**d**, A 100× dark field image showing the 2D optical grating pattern on individual micromirrors.

Type or paste caption here. Create a page break and paste in the figure above the caption.

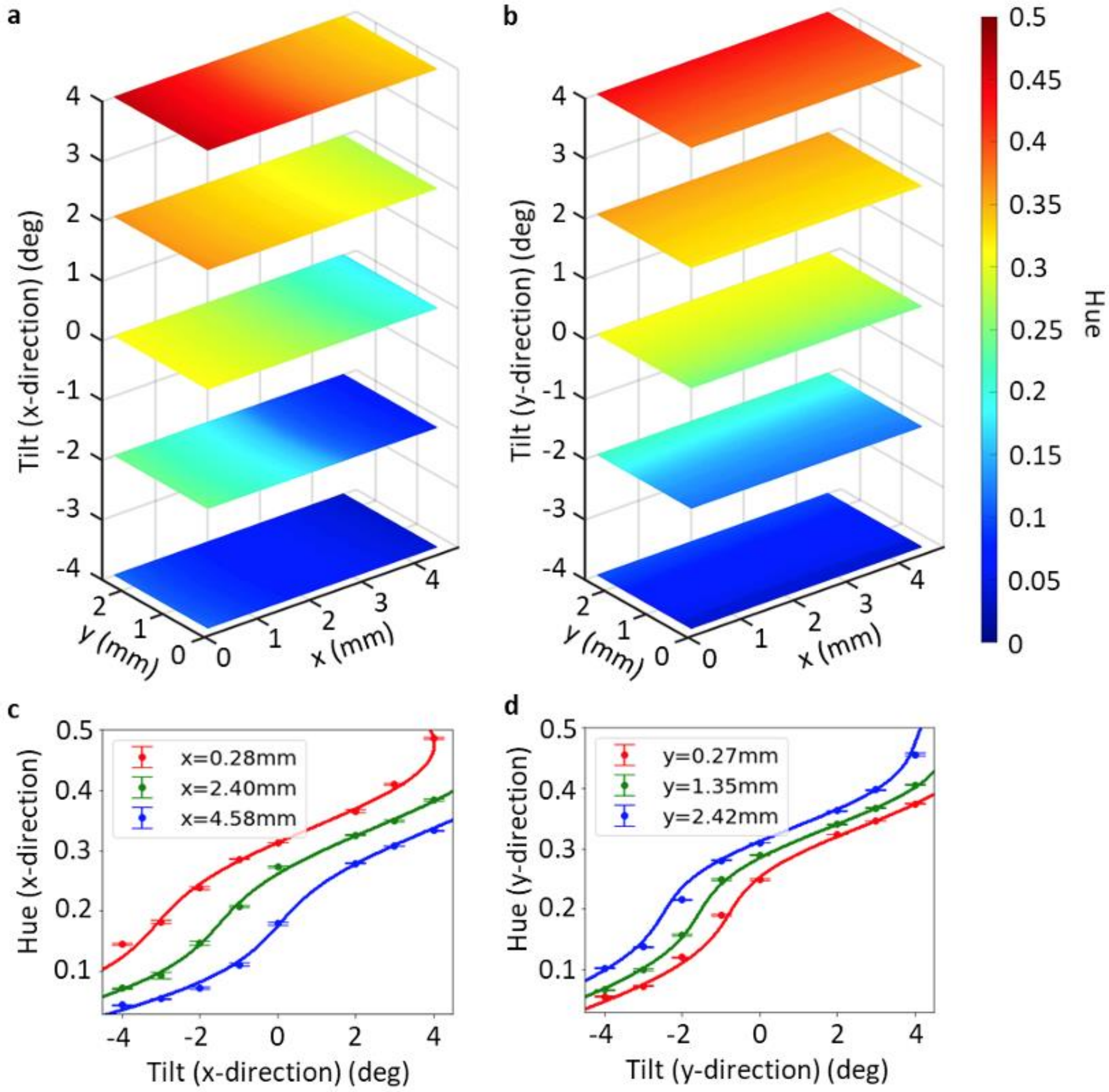


**Fig. S2.**

**Calibrating relationships between tilting angles, hue, and position using multi-dimensional curve fitting.**

**a**, The tilting angle in the x-direction correlated with x and y spatial positions of a micromirror and its hue value. For micromirrors of the same tilting angle, there is a gradual change of hue along the x axis, and negligible change along the y axis. We sampled experimental data at nine positions, around the edge and the center of the whole FOV on the SPOT microscopy platform.

**b**, For tilting angles in the y-direction, hue value changes gradually along the y axis, and negligibly along the x axis.

**c**, The relationship between the hue (x-direction) and micromirror tilting angle in the x-direction is non-linear. Radial basis functions were used to optimize fit for this relationship.

**d**, The relationship between the hue (y-direction) and tilting angle in the y-direction.

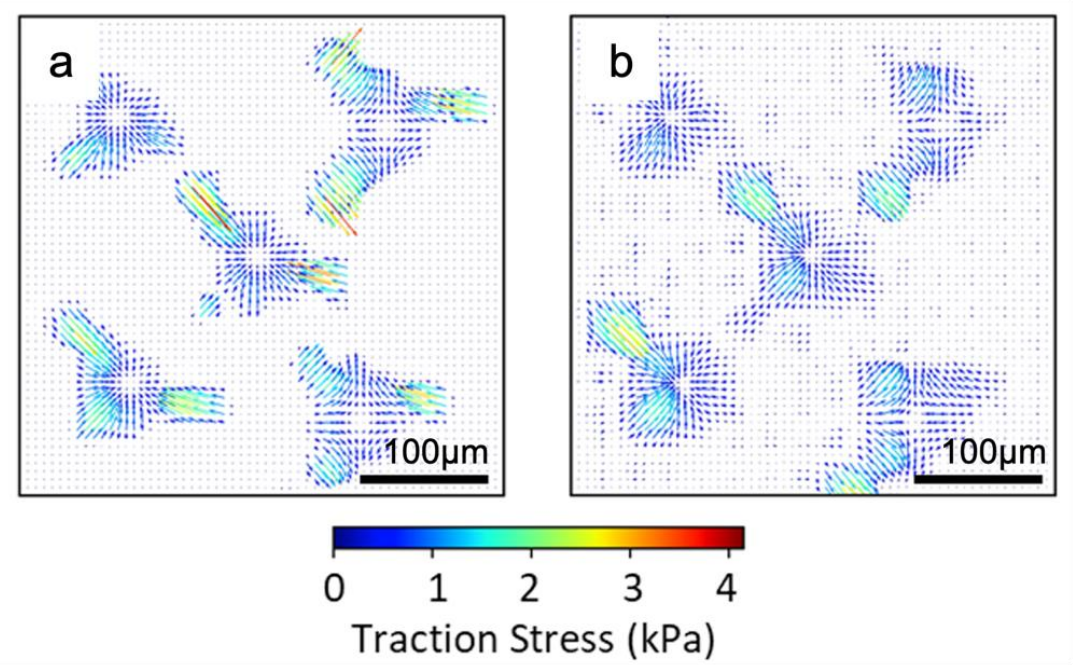


**Fig. S3.**

**Spatial resolution study of traction force distribution measured by SPOT microscopy.**
**a,** A known experimental traction force distribution extracted from a prior study (*31*) measuring the contraction force of C2C12 myocytes. This known distribution was imported into the COMSOL model to solve for the corresponding 2D micromirror tilting angles.
**b,** The tilting angles corresponding to (**a**) were fed into our trained machine learning model to predict the traction force distribution. The positions and shapes of the predicted traction force distribution matches well with the known traction force distribution in (**a**). The predicted model in (**b**) has a smoother traction force distribution because of the use of a Fourier series of limited spatial frequencies. These results in (**b**) show a strong concordance with the sub-cellular traction force distribution in (**a**).

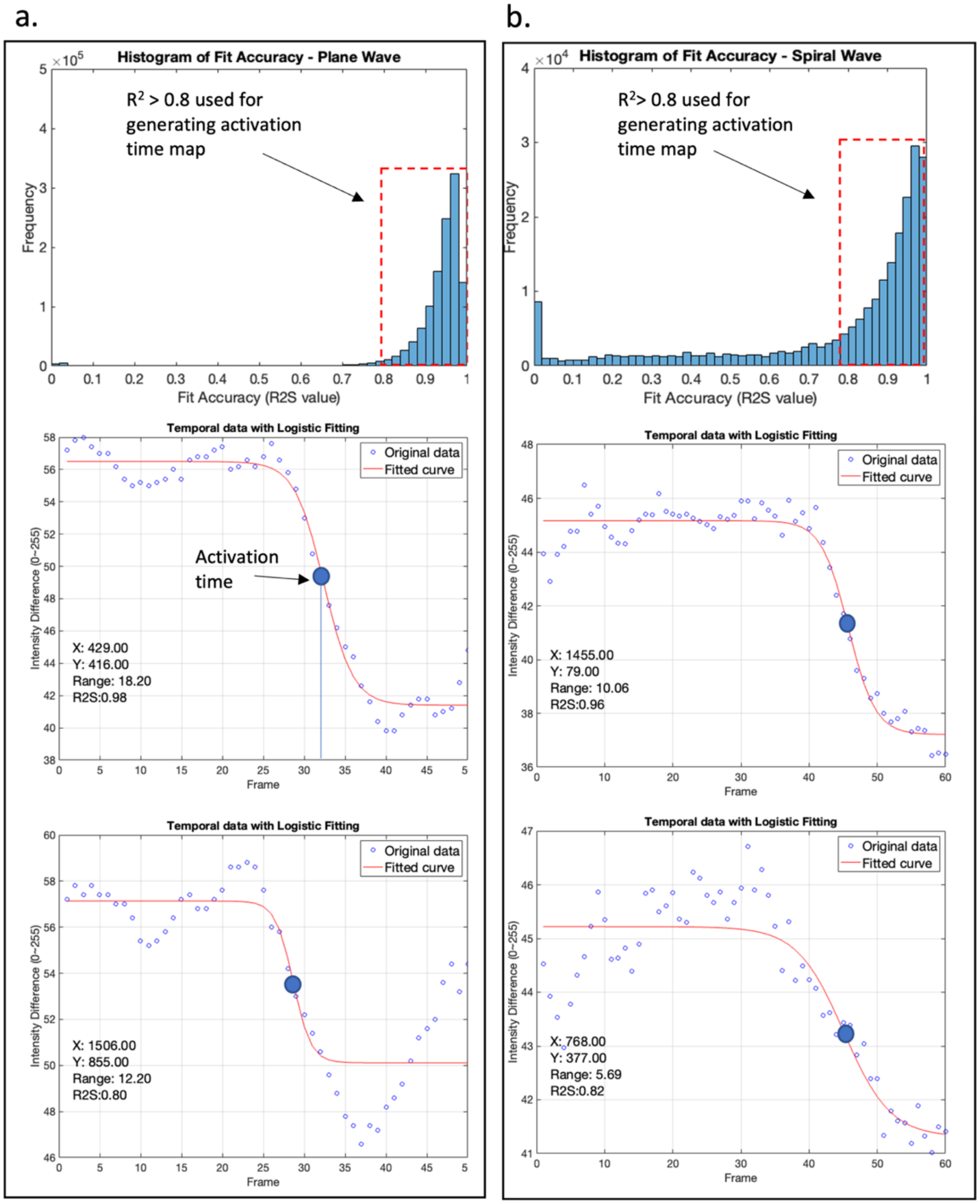


**Fig. S4.**

**Extraction of activation times from temporal intensity profiles.**

The extraction of the activation time of each pixel is by curve fitting using the logistic function. The activation time is determined at the point where the slope of the fitting curve reaches its maximum.

**a,** Histogram of fitting accuracy for NRVM propagation of a plane mechanical wave. Only pixels with coefficient of determination ($R^2$) larger than 0.8 (1 represents perfect fitting) are chosen for constructing the activation time map shown in Fig. 3a. Example temporal intensity profiles and their corresponding fitting curves with $R^2$ between 0.8~1 are presented for reference.

**b**, Histogram and example temporal data for NVRM propagation of a spiral mechanical wave.

**Movie S1.**

Detection of mechanical wave propagation on SPOT without using fluorescence dyes. Processed SPOT video showing coordinated cardiomyocytes beating across a cm scale in less than a second. The field-of-view is 9 mm × 9 mm. The color represents the green pixel intensity change. The video is played in real-time at 44 frames per second. The RGB colors measured in the SPOT images contain both the static and dynamic components. By subtracting the static term from the original image, the dynamic color component that demonstrates the dynamic contraction and relaxation behaviors of mechanical wave propagating across a tissue can be extracted.

**Movie S2.**

A spiral wave of NRVM tissue beating captured on SPOT with a substrate of stiffness 21 kPa. The field-of-view is 4.82 mm × 2.71 mm. The color represents the green pixel intensity change. The real-time frame rate captured by the high-speed camera is 83 frames per second. The video frames were time-averaged over 5 frames to reduce the noise. The mechanical wave initiates from the center and progresses in the counterclockwise direction.
